# Human lung organoids develop into adult airway-like structures directed by physico-chemical biomaterial properties

**DOI:** 10.1101/564252

**Authors:** Briana R. Dye, Richard L. Youngblood, Robert S. Oakes, Tadas Kasputis, Daniel W. Clough, Melinda S. Nagy, Jason R. Spence, Lonnie D. Shea

## Abstract

Tissues derived from human pluripotent stem cell (hPSC) often represent early developmental time points. Yet, when transplanted into immunocompromised mice, these hPSC-derived tissues further mature, which has been enhanced with biomaterial scaffolds, gaining tissue structure and cell types similar to the native adult lung. Our goal was to define the physico-chemical biomaterial properties, including the polymer type, degradation, and pore interconnectivity of the scaffolds. Transplantation of human lung organoids (HLOs) on microporous poly(lactide-co-glycolide) (PLG) scaffolds or polycaprolactone (PCL) produced organoids that formed tube-like structures that resembled both the structure and cellular diversity of an adult lung airway. Microporous scaffolds formed from poly(ethylene glycol) (PEG) hydrogel scaffolds inhibit maturation and the HLOs remain as lung progenitors. The structures formed from cells that occupy multiple pores within the scaffold, and pore interconnectivity and polymer degradation contributed to the maturation. Finally, the overall size of the generated airway structure and the total size of the tissue was influenced by the material degradation rate. Collectively, these biomaterial platforms provide a set of tools to promote maturation of the tissues and to control the size and structure of the organoids.

## Introduction

Human lung organoid models allow for the study of cell fate decision during development in addition to modeling diseases such as cystic fibrosis and goblet cell metaplasia, along with infections such as respiratory syncytial virus (Chen et al., 2017; Danahay et al., 2015; Dye et al., 2015; McCauley et al., 2017; Miller et al., 2017; Rock et al., 2009; Tadokoro et al., 2014). Organoids represent aspects of the native human organ including structure and cellular organization. The properties of the organoid enable studies on human development, tissue regeneration and disease, for which we have not previously been able to perform (Dedhia, Bertaux-Skeirik, Zavros, & Spence, 2016; Dye, Miller, & Spence, 2016b; Fatehullah, Tan, & Barker, 2016; Huch & Koo, 2015; Johnson & Hockemeyer, 2015; Rookmaaker, Schutgens, Verhaar, & Clevers, 2015). In particular, organoids have been employed to model lung airways *in vitro* with microfluidic platforms, in which an air-liquid interface is employed to recreate the microenvironment of the airway epithelium (Gkatzis, Taghizadeh, Huh, Stainier, & Bellusci, 2018; Miller & Spence, 2017). These platforms are amenable to high-throughput screening and studying specific cellular response for drug testing and toxicity in short term culture. We have previously derived human lung organoids that incorporates tissue structure including both the epithelium and supporting tissue (cartilage, smooth muscle, fibroblasts) within long term cultures (Dye et al., 2015; 2016a). This development of lung structures enable the study of airway diseases that affect either the epithelium such as chronic obstructive pulmonary disease (COPD) or the supporting tissue such as asthma (Athanazio, 2012). Notably, these *in vitro* cultures to date generate structures that reflect the fetal airway.

Adult airway structures can be generated with HLOs through *in vivo* transplantation (Dye et al., 2016a). This observation with *in vivo* transplantation of HLOs reflects observations with numerous organoid systems (Camp et al., 2015; Dye et al., 2015; Finkbeiner et al., 2015; Takasato et al., 2015), that the in vivo environment supports maturation to adult tissue phenotypes and structures (Dye et al., 2016a; Finkbeiner et al., 2015). We had previously derived human lung organoids (HLOs) from pluripotent stem cells (hPSCs) by an initial *in vitro* culture that targets signaling pathways during lung development. After deriving definitive endoderm, the 3D structures formed from the monolayer and express the early lung marker NKX2.1 and anterior foregut marker SOX2. These early 3D structures called foregut spheroids were placed into a Matrigel droplet overlaid with media supplemented with FGF10. The foregut spheroids then grew into larger organoids, with the transcriptome analysis indicating that HLO represented fetal lung (Dye et al., 2015). In order to surpass this developmental barrier, lung organoids were transplanted onto a microporous polymer scaffold into the epididymal fat pad of immunocompromised mice. After 8 weeks, the transplanted HLO (tHLO) had airway-like structures indistinguishable from native adult airways including proper cellular organization, epithelial cellular ratios, airway cell types, and surrounded by smooth muscle and cartilage (Dye et al., 2016a). Interestingly, this maturation to adult airway structures occurred on the microporous scaffolds, yet not in more typical transplantation platforms (Matrigel droplets into either the kidney capsule, omentum, or epididymal fat pads).

In this report, we investigated the physico-chemcial properties of microporous scaffolds that support HLO maturation into airway structures. Microporous scaffolds were formed from multiple materials using poly(lactide-co-glycolide)(PLG), polycaprolactone (PCL), and poly(ethylene glycol) (PEG). The polymers may have distinct interactions with the host, and also the pore interconnectedness was varied through the initial fabrication and also through the degradation rate of the polymers. For these material platforms, we investigated airway maturation, immune response, as well as overall explant and airway size. Identifying the biomaterial design parameters that influence airway maturation and structure will enable the development of platforms that can direct the structure to better model airway homeostasis and disease environments.

## Results

### PEG, PCL, and PLG have varying extent of HLO derived airway maturation

Microporous scaffolds with similar architectures and formed from 75:25 (lactide:glycolide) PLG and 20% (w/v) 4-arm PEG-maleimide microporous scaffold were seeded with foregut spheroids. Both scaffolds had pores ranging in size from 275μm to 425μm, and were cylinders having a diameter of 5mm diameter and a thickness of 2mm. Note that PLG is a degradable, hydrophobic polyester that will adsorb proteins to support HLO adhesion. The PEG scaffolds are non-degradable hydrogels and were formed with or without the adhesion peptide RGD, which would support adhesion of the HLOs to the hydrogel. The seeded PLG and PEG scaffolds were cultured for 7 days *in vitro*, during which time the foregut spheroids grow to fill the pores (Figure 1A). Scaffolds were then transplanted and retrieved after 8 weeks. For the PLG scaffolds, tissue was found within and around the scaffold, and the tissue throughout contained airway-like structures. The majority of the PLG scaffold degraded and the material was not detected in histological sections. The PLG tHLO had organized into pseudostratified epithelium resembling native adult airways (Figure 1C).

**Figure 1.**
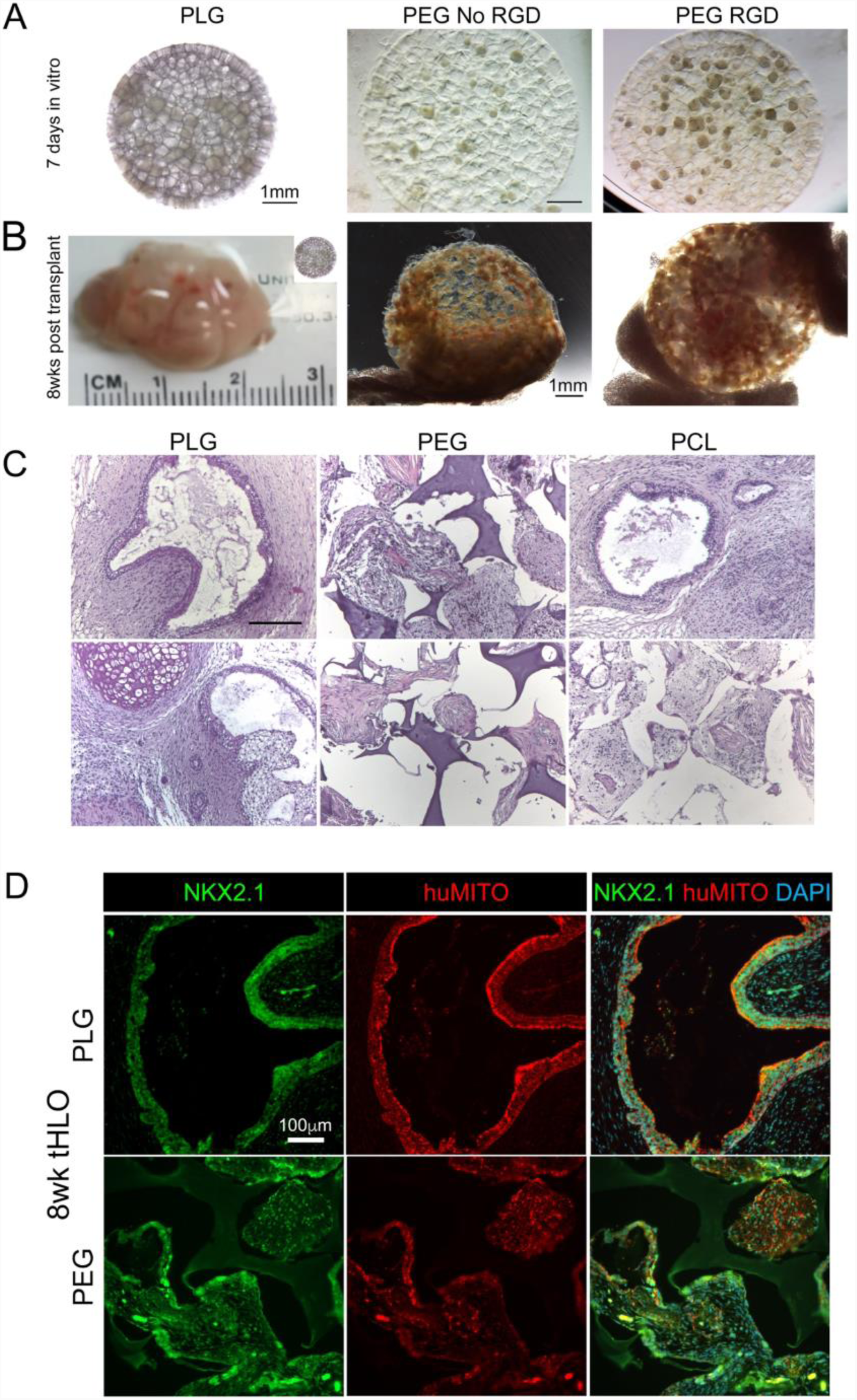
HLOs seeded on PEG, PLG, and PCL scaffolds affects airway structure formation. A) Approximately 50 foregut spheroids were seeded onto PLG and PEG with or without RGD. Wholemount images were taken after 1 week in culture. B) Scaffolds were transplanted into the epididymal fat pad of immunocompromised mice and retrieved 8 weeks later. The PLG tHLO grew up to 2.5cm while the tissue remained within the PEG scaffolds with or without RGD. C) The histology of the PLG and PCL tHLOs had organized pseudostratified epithelium resembling native airway epithelium. The PEG tHLOs and the tissue within the pores of the PCL had no organized epithelium, but remained as clusters of cells. D) Some of the clusters of cells within the PEG scaffold were lung marker NKX2.1+ and human mitochondria (huMITO)+. Scalebars for A-B:1mm, C:200μm, and D:100μm.

This growth within and around the scaffold observed with PLG contrasted with the PEG tHLO, which remained within the pores of the PEG scaffold independent of whether the scaffold was modified with or without RGD modification (Figure 1B). We hypothesized that the individual HLOs would grow together on the surface of the PEG scaffold and form airway-like structures. However, the PEG tHLOs remained within separate pores. We investigated the possibility that smaller airways formed within the pores of the PEG scaffold. However, the PEG tHLO only had clusters of cells within the pores with no organized, pseodostatified epithelium that resemble native airway structures compared to the PLG tHLO, which possessed pseudostratified epithelial structures report (Figure 1C)(Dye et al., 2016a). The tissue within the PEG pores was derived from the HLOs since the cells expressed human mitochondria maker (huMITO) in addition to the host cells filling the pores (huMITO-cells). The huMITO+ cells expressed lung marker NKX2.1, yet no mature cell types or airway-like structures were observed in the PEG tHLO explants (Figure 1D). In comparison, the organized airway structures observed in the PLG tHLO explants expressed lung marker NKX2.1 and were derived from the hPSC since the structures were huMITO+.

PCL scaffolds were subsequently transplanted with HLOs, which was motivated by having a hydrophobic polyester polymer with a slower degradation rate than the PLG, yet a similar degradation as PEG. Following HLO seeding on PCL scaffolds and transplantation for 8 weeks, the explants contained organized, pseudostratified airway structures similar to the structures observed in the PLG tHLOs. Yet, the structures in the PCL tHLOs surrounded the scaffold and no organized epithelial structures were observed within the pores of the PCL but instead had clusters of cells similar to what is observed in the PEG tHLO. The PCL scaffold was intact similar to the PEG, and the tissue surrounding the scaffold contained airway-like structures, while the tissue within the pores resembled the PEG tHLOs (Figure 1C). Collectively, the polyester polymers supported the development of airway structures after 8 weeks, and while the microporous architecture supports seeding of the HLOs, the structures do not form within intact pores of the scaffold.

### Initial immune response at microporous scaffolds may contribute to HLO responses

The initial immune response within the microporous scaffolds was investigated as a potential mechanism underlying the differential maturation on the various materials. Lung organoids were transplanted and collected after 1 week *in vivo*, with analysis of the innate immune response. Overall, PEG tHLOs had a significantly greater percentage of leukocytes (CD45+) than PLG and PCL tHLOs (Figure 2A), which indicates greater cell recruitment from the host tissue. From the CD45+ population, the PEG tHLOs had significantly higher percent of CD11b+GR1+Ly6c-cells, often referred to as myeloid derived suppressor cells, and significantly less CD11c+F4-80- (dendritic) cells and Ly6c+F4-80- (monocyte) cells compared to PLG and PCL tHLOs (Figure 2B). No differences were observed between the PLG and PCL scaffolds. These data suggest that the immune response could be a contributing factor to the inhibition of HLO maturation in PEG scaffolds relative to PLG and PCL scaffolds.

**Figure 2.**
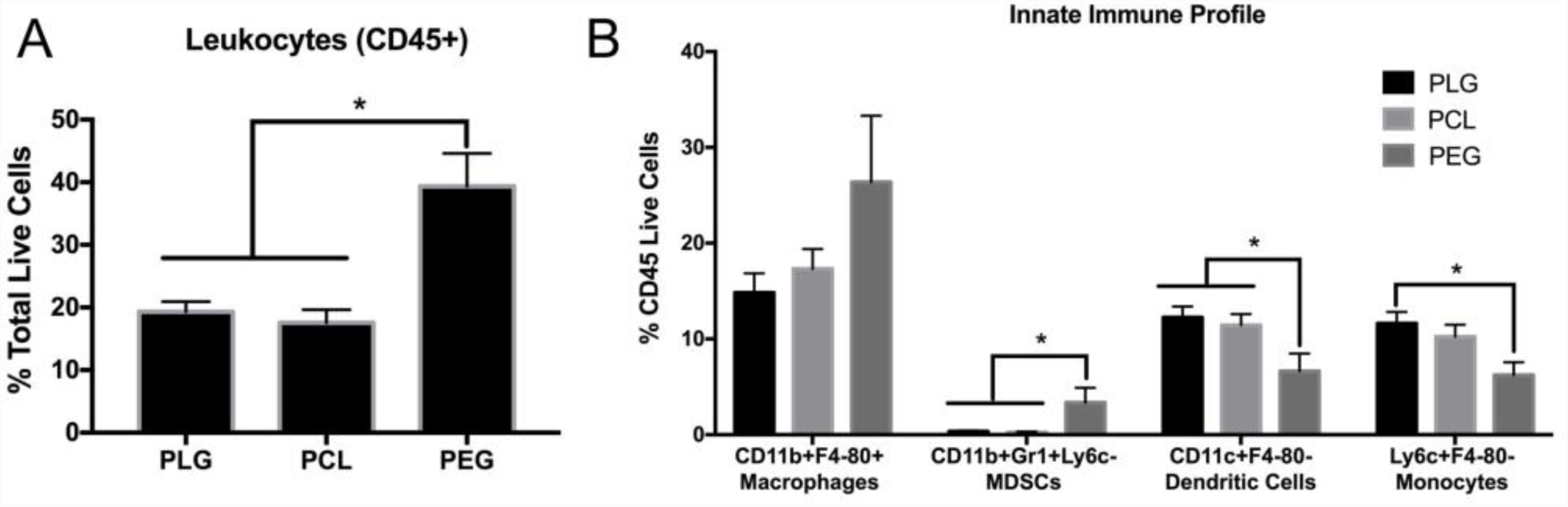
The innate immune profile for PEG, PLG, and PCL tHLOs. Foregut spheroids were cultured on scaffolds for 1 week *in vitro*, transplanted into the mouse epdidymal fat pad, and then retrieved after 1 week to observe the innate immune response. A) The PEG tHLOs had 39% CD45+ Leukocytes (N=6) compared to 17% CD45+ cells in PLG tHLOs (N=7) and 19% CD45+ cells in PCL tHLOs (N=7) P<.005 B) PEG, PCL and PLG and similar percent of macrophages (CD11b+F4-80+). PEG tHLOs had significantly higher MDSCs (CD11b+GR1+Ly6c-) compared to PLG and PCL tHLOs, *P<.05. In contrast, PEG tHLO had significantly lower dendritic cells (CD11c+F4-80-) and monocytes (Ly6c+F480-) *P<.05. All error bars represent SEM.

### HLO fusion during formation of airway-like structures

We subsequently investigated the contribution of HLO interaction in adjacent pores to the formation of airway structures, which was motivated by the observations of airway structures forming on the surface of slow degrading PCL scaffolds. We analyzed for fusion of multiple organoids into an airway structure by seeding scaffolds with GFP+ HLOs and GFP-HLOs. Following culture of HLOs in PLG and PCL scaffolds for 1 week *in vitro*, we observed pores with GFP+ HLOs were adjacent to pores containing GFP-HLOs (Figure 3A). Scaffold were transplanted and retrieved after 4 and 8 weeks *in vivo*. After 4 weeks *in vivo*, airway structures were not present, yet there were populations expressing lung marker NKX2.1 that were either GFP+ or GFP-indicating both HLO populations survived and successfully generated lung progenitors (Figure 3B). After 8 weeks, airway structures were observed in PLG and PCL tHLOs, and these structures contained lung airway ciliated cells (ACTTUB+) and basal cells (P63+). These airway structures had both GFP+ and GFP-cells both in PCL and PLG tHLOs (Figure 3C), indicating that the airway structures form by the HLOs fusing from adjacent pores in both PCL and 75:25 PLG tHLOs.

**Figure 3.**
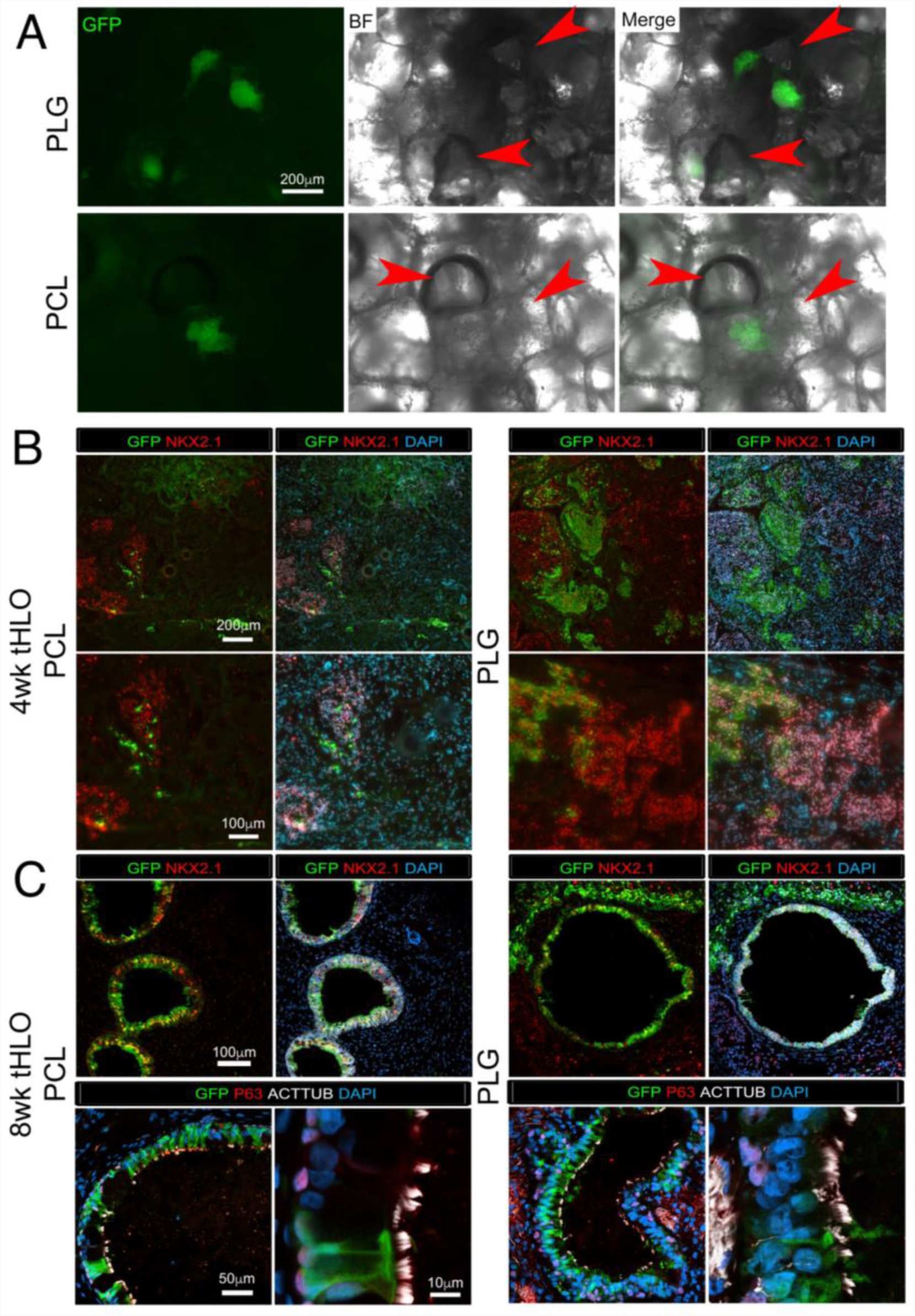
The degradation rate of the scaffold affected airway diameter. A) The average measurement was taken of the longest and shortest diameter for each cross section of an airway structure in a 8wk tHLO. The 85:15 PLG tHLOs (224μm) had the significantly shorter diameter compared to the 75:25 control PLG (333μm) *P<.05. The 50:50 PLG tHLO had the longest diameter at 415μm. The PCL control and large interconnected PCL tHLOs had similar diameter at 277μm and 299μm respectively compared to the 85:15 PLG tHLO (224μm). All error bars represent SEM. B-C) Histology sections of PLG (75:25), large interconnected PCL, 85:15 PLG and 50:50 PLG represent the quantified sections. Scale bars B) 200μm and C) 400μm.

We next tested the hypothesis that an increase the size of pore interconnections would increase HLO fusion and create larger airway structure. We obtained PCL scaffolds constructed through 3D printing rods of PCL in a cross-hatch pattern within a 5mm wide, 2mm tall cylinder (same dimensions as the scaffold fabricated in lab), which has large pore connections (300μm) relative to the PCL scaffolds (ranging from 10 – 100μm)(Rao et al., 2016). The size of the airway structures was similar between both types of PCL scaffolds (Figure 4A, B). These results suggest that while fusion of cells in adjacent pores contributes to the formation of airway structures, yet other factors are determining the size of the structures.

**Figure 4.**
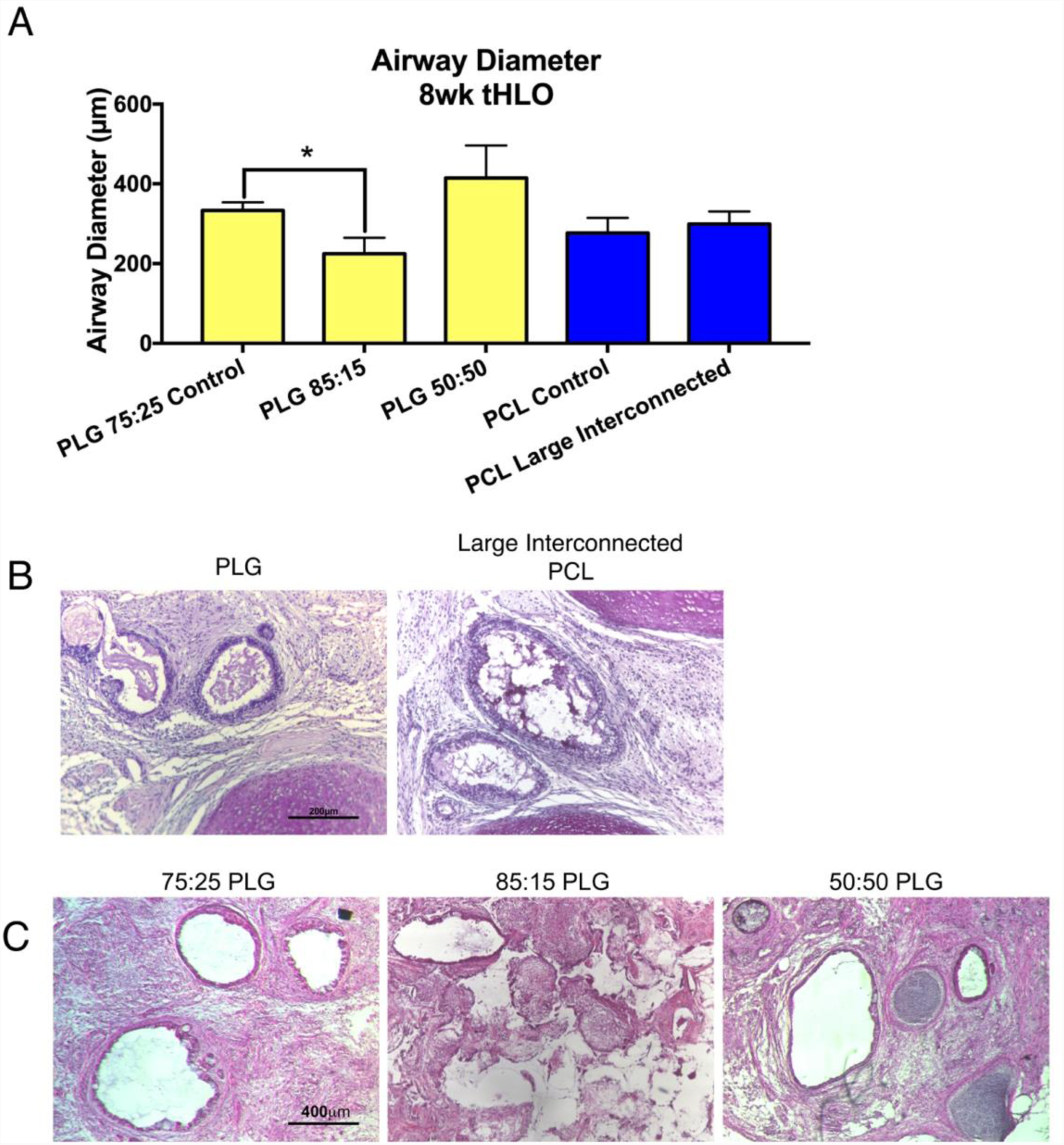
The degradation rate of the scaffold affected explant size of tHLO. A) Overall, the 75:25 PLG had the largest explant size (1.18cm, N=6) after an 8wk transplant compared to both fast (50:50 PLG, N=5) and slow degrading (85:15 and PCL, N=5) *P<.05. The fast degrading 50:50 PLG tHLO was significantly larger than the 85:15 PLG tHLO explant *P<.005. B) Ki67+ cells (red) were present within the epithelial airway structures labelled with ECAD (green) and the surrounding tissue both in PCL and 75:15 PLG. C) 75:25 PLG (19.6% ±1.8%) had significantly more Ki67+ cells than PCL (8.7% ±1.8%). D) The 75:25 PLG had significantly more Ki67+ cells in the ECAD+ and ECAD-areas (ECAD+: 6.7 ±0.9, ECAD-: 13.8% 1.7) compared to PCL tHLO (ECAD+: 3.009 ±1.0, ECAD-: 6.4% 1.1). Both for PCL and PLG tHLOs there was significantly more Ki67+ cells in the surrounding tissue (ECAD-) compared to the organized epithelium (ECAD+). **P>.005, *P>.05 All error bars represent SEM.

### Scaffold degradation affects the HLO derived airway size

Given the fusion of HLOs in adjacent pores, we next investigated the contribution of polymer degradation to the number of airway-like structures, which could ultimately affect the size of the airway. The airway diameters in the PCL tHLOs trended towards being slightly smaller (276μm) compared to 75:25 PLG tHLOs (333μm, Figure 4A). We also transplanted HLOs onto 85:15 PLG polymer which degrades at a rate intermediate of that between 75:25 PLG and PCL. The 85:15 PLG tHLOs had significantly smaller airway diameter (224μm, p=0.049) than the 75:25 PLG (Figure 4C). In addition, the 85:15 PLG tHLO had a similar phenotype to the PCL tHLO, with the airway-like structures present adjacent to the scaffold and the tissue within the pores remaining as cell clusters (Figure 3C). We also investigated a faster degrading polymer, 50:50 PLG, with the hypothesis that it would support larger airway-like structures. The 50:50 tHLOs trended towards larger airways (414μm) relative to the 75:25 PLG (Figure 4C), though the difference not significant. Collectively, the size of the airway structures was influenced by the degradation rate of the scaffold, with slower degrading polymers leading to smaller airway-like structures.

### Controlling the tHLO explant size

The size of the explant varied as a function of the polymer. The size of the explant for HLOs transplanted on PLG scaffold reached diameters up to 2.5 cm, which is five times the original scaffold diameter (5mm). The PCL tHLO explant size was significantly smaller than 75:25 PLG tHLOs, with the PCL explants having diameters in the range of 0.5 to 0.6 cm (Figure 1A, 5A). The explant sizes in fast degrading 50:50 PLG, slow degrading 85:15, and large interconnected PCL scaffolds were 0.81 cm, 0.53 cm, and 0.65 cm respectively. Relative to 75:25 PLG tHLOs, the other tHLO conditions had a significant reduction in explant size. However, the 50:50 PLG tHLOs were significantly larger than the 85:15 PLG tHLO. All together, these data suggested that both the slow and fast degrading PLG caused a reduction in explant size, but the size reduction was more significant in the slow degrading polymers, 85:15 PLG and PCL. Both the PCL control and PCL large interconnected pores were similar in size (Figure 5A). Thus, the size of pore interconnections does not appear to significantly impact the explant size.

**Figure 5.**
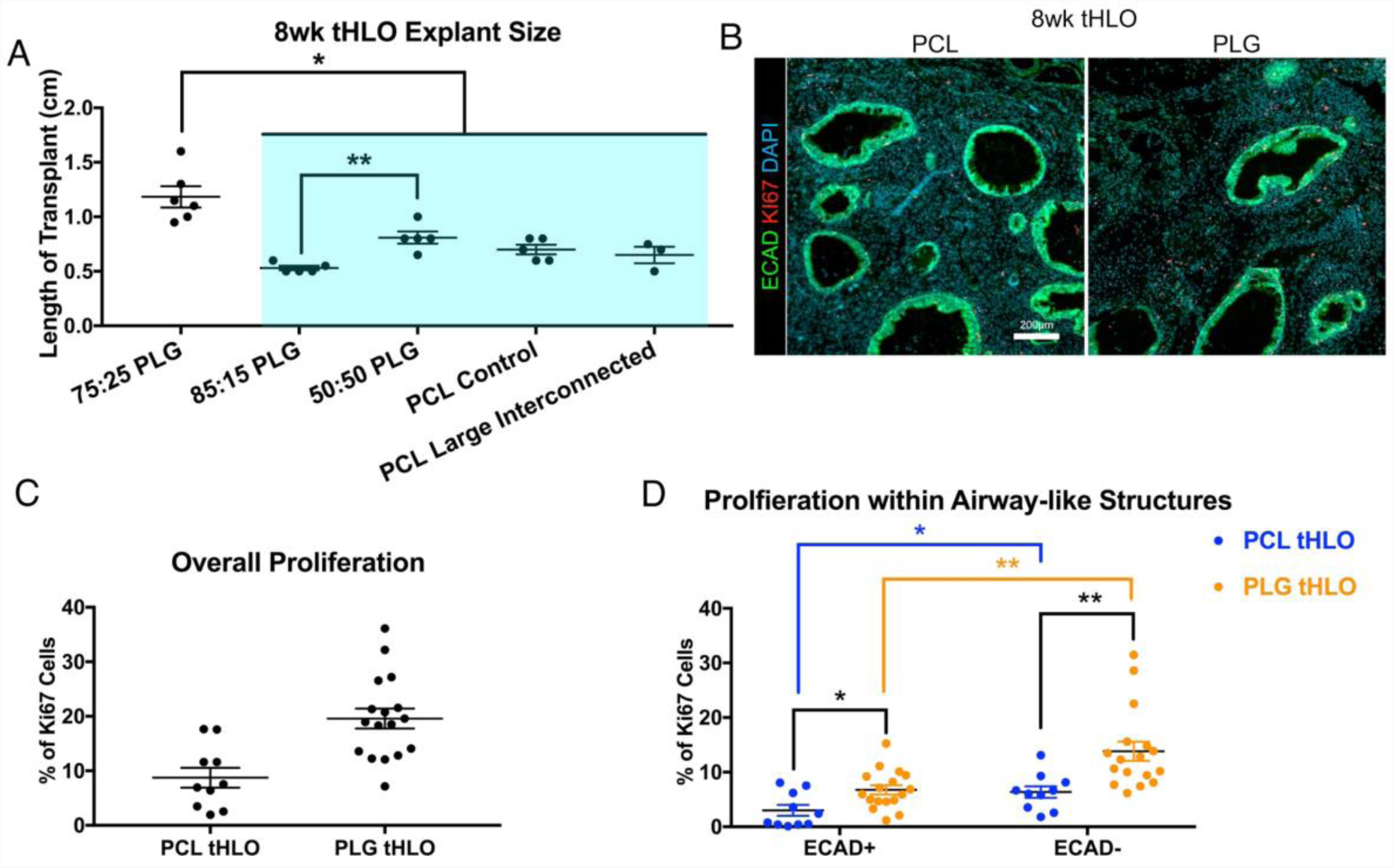
HLOs fuse together to form airway structures. A) GFP H1 hESC line and H1 hESC derived HLOs were seeded onto 75:25 PLG and PCL scaffolds. Wholemount images of the scaffolds cultured for 1 week *in vitro* had GFP+ HLOs next to a pore of GFP-HLOs indicated by the red arrow head in both PLG and PCL conditions. B) After transplanted for four weeks, the GFP+ cells were mixing with the GFP-cells and both expressed early lung marker NKX2.1 (red). No organized epithelial structures were observed for either scaffold. C) After transplanted for 8 weeks, airway structures formed and were comprised of GFP+ and GFP-cells in both PLG and PCL tHLOs. The airway structures expressed NKX2.1 (red). D) The fused GFP+ and GFP-HLOs in both PLG and PCL had multiciliated cells labelled by acetylated tubulin (ACTTUB, white) and basal cell marker P63 (red). Scale bars represent A-B 200μm, B-C 100μm, and C-D 50μm, 10μm.

Proliferation within the explant was subsequently investigated as a contributing factor to the explant size. A significant two-fold change in proliferation (Ki67+cells) in 75:25 PLG tHLO (19.5%) was observed relative to PCL tHLO (8.7%) (Figure 5B-C). This increase in proliferation was observed both in the airway-like structures (ECAD+) and the surrounding tissue (ECAD-). Interestingly, both in PLG and PCL tHLOs, proliferation was significantly greater in the tissue adjacent to the scaffold relative to the ECAD+ airway-like structures (Figure 5D).

## Discussion

In this report, we have demonstrated that the type of material and degradation of the microporous scaffold can affect lung airway formation, airway size, and explant size derived from transplanted HLOs. Previously, 75:25 PLG microporous scaffolds were used to transplant HLOs into the epididymal fat pad (Dye et al., 2016a). We hypothesized that the tHLOs needed a surface to grow and expand on in order to mature in airway structures, since no maturation occurred when HLOs were placed into the kidney capsule or sewn in the omentum of an immunocompromised mouse. We then tested this hypothesis and found that the HLOs did not require the support of the pores to form airway-like structures, but instead needed specific material properties to allow for the development of airway structures. More specifically, the degradation of the material affected airway size and overall explant size of the tHLO.

As the airway structures formed, the individual HLOs fused together to form these structures evident by the GFP+ and GFP-HLOs forming into one airway. Interestingly, the buds forming together occurred when the scaffold held its shape (PCL) or degraded (75:25 PLG) over the 8 weeks *in vivo*. Thus, the degradation was not necessary for the HLOs to fuse together to form the airway structures; however, there were no structures within the pores of the scaffolds that held the scaffold structure during the 8 weeks (PCL and 85:15 PLG). The fusion of the HLOs took place either on the surface of the scaffold or as the scaffold degraded. The large pore interconnected PCL allowed for airways to form within the scaffold. Collectively, the HLOs could only form airway structures if they had the space to fuse and expand together into epithelial tubes. This need of expansion aligns with airway formation during lung development where the airways start as one epithelial bud that continually bifurcates and expands into the surrounding mesenchyme to ultimately form a network of airway (Metzger, Klein, Martin, & Krasnow, 2008; Rawlins, 2010).

The degradation of the material contributed to multiple properties of the organoids: airway size and overall explant size. Deriving variations in airway size will allow the study of airway diseases such as COPD and asthma in both large and small airway models (Athanazio, 2012). An airway ranging in size from 200-350μm represents a 5^th^ generation airway in a native human lung while the 4^th^ generation ranges from 400-600μm (Lewis et al., 2005). Here by changing the degradation we were able to represent two types of airways, 4^th^ and 5^th^ generation size that are observed in the native adult lung. The data with PLG and PCL indicated that the airway size was maximal for 75:25 PLG, with slightly smaller structures for 50:50 PLG and 85:15 PLG, suggesting that degradation plays a role and that maximal size occurs at an intermediate rate of degradation. One mechanism by which degradation can influence airway size is through the fusion of organoids from adjacent pores. Polymer degradation would influence fusion by increasing the size of the interconnections between pores over time, which would allow for greater connectivity. Polymer degradation would also function to remove the polymer as a substrate for organoid development, which may also influence growth and maturation. Collectively, these results are consistent with the general idea that the polymer degradation should be matched with the rate of tissue formation.

Multiple scaffolds maintained the ability to form airway structures, yet the explant size was a function of the scaffold properties. PCL scaffolds allowed for the formation of structures at the surface of the material, yet not within the pores of the scaffold. The increased size of the explant did not influence the size of the airway-like structures that formed, and the increased size resulted from an increase of proliferation from the supporting tissue including the mesenchyme with a lesser extent increase of proliferation from the organized epithelium. The scaffold properties influenced the proliferation of the progenitor cells, that subsequently influenced the overall size of the explant. A more controlled growth of the organoid would allow for longer studies to be done, since the explant growth will not impede the mouse health. For instance, HLOs could be generated from patient specific hPSCs lines that have Cystic Fibrosis (CF). Patients with CF have mutations in Cystic Fibrosis Transmembrane conductance Regulator (CFTR), which causes excess of mucus within the airways which leads to chronic infection and inflammation of the lung epithelium(Cutting, 2015). With this model, HLOs could be generated from CF patient specific hPCS and studied in the tHLO model that provides a human airway model. Since the explant size is smaller, longer studies can be conducted in order to understand the short and long-term effects of CF on the airway epithelium and surrounding tissue including smooth muscle, cartilage and vasculature.

Microporous scaffolds compose of PEG did not support maturation over the 8 weeks and the HLOs remained as NKX2.1 progenitors. HLOs were seeded onto PEG hydrogels with and without modification with RGD, a fibronectin binding peptide, in order to investigate if ECM signaling may be a signal directing maturation. The presence of RGD peptide did not impact maturation, suggesting that either adhesion is not a limiting factor in HLO maturation or that the RGD is insufficient to trigger the necessary signaling cascades. The innate immune response, which is active in immunocompromised mice, differed significantly between PEG versus PLG and PCL. This difference could be a contributing factor that influenced HLOs maturation since it is known that the immune system affects hPSCs and tissue regeneration including adult stem cells (Aurora & Olson, 2014; Kizil, Kyritsis, & Brand, 2015; Pearl, Kean, Davis, & Wu, 2012).

Overall, HLO maturation was supported by multiple microporous scaffolds that resulted from fusion of organoid clusters in adjacent pores. The physico-chemical properties of the scaffold influenced the properties of explant, such as the number and size of airways structures and the size of the explant. The biomaterials thus provide a tool that may be capable of directing tissue formation from organoids for the purpose of modeling normal development, and also for modeling disease states. Specific to airways, controlling airway and total explant size will allow for new models for airway diseases such as asthma, COPD, and CF with the potential to perform long-term studies.

## Methods

### Maintenance of hESCs, generation of HLOs, and seeding on scaffolds

H1 human embryonic stem cell (hESC) line (NIH registry #0043) and H9 (NIH registry #0062) was obtained from the WiCell Research Institute. H1 hESC line was used to derive all HLOs for these experiments except for Figure 2 where H9 hESC and H9 GFP hESC lines were used to derive HLOs. H9 GFP hESC line was generated by infecting H9 hESCs with pLenti PGK GFP Puro virus generated from the plasmid purchased from AddGene (Cat#: 19070)(Campeau et al., 2009). H9 GFP hESC clonal line was generated by puromycin selection flow cytometry analysis sorting (FACS) for GFP high expressing cells. All hESC lines were approved by the University of Michigan Human Pluripotent Stem Cell Research Oversight Committee. hESCs were maintained as previously described (Spence et al., 2011). HLOs were derived as previously described (Dye et al., 2015). Foregut spheroids, which grow into HLOs, were seeded on scaffolds as previously described (Dye et al., 2016a).

### Scaffold fabrication

75:25 PLG used and scaffolds were fabricated as previously described (Blomeier et al., 2006; Dye et al., 2016a). 85:15 (Resomer® RG 858 S, Poly(D,L-lactide-co-glycolide), Sigma, Cat#: 739979-1G) and 50:50 PLG (Resomer® RG 505, Poly(D,L-lactide-co-glycolide) ester terminated, MW: 54,000-69,000, Sigma,Cat#: 739960) were fabricated the same as 75:25 PLG scaffolds. 20% 4arm PEG malimide, 20,000MW (JenChem, Cat#: A7029-1) hydrogels were fabricated as previously described(Rios et al., 2018). PCL scaffolds were fabricated as previously described(Rao et al., 2016). Large interconnected PCL were commercially bought (National Institute of Standards and Technology, Cat#:8394).

### Immunohistochemistry, hematoxylin and eosin stain (H&E), and imaging

Immunostaining and H&E were carried out as previously described (Rockich et al., 2013). Antibody information and dilutions can be found in Table 1. All images and videos were taken on a Nikon A1 confocal microscope or the Zeiss Axio Observer.Z1.

### Flow Cytometry

Cell disassociation and flow cytometry was previously described(Rao et al., 2016). Antibodies used are as follow: Fluor® 700 anti-CD45 (1:125, clone 30-F11, Biolegend), V500 anti-CD11b (1:100, clone M1/70, BD Biosciences), FITC anti-Ly6C (1:100, clone HK.14, Biolegend), PE-Cy7 anti-F4/80 (1:80, clone BM8, Biolegend), APC anti-CD11c (1:80, clone N418, Biolegend), and Pacific Blue™ anti-Ly-6G/Ly-6C (Gr-1) (1:70, clone RB6-8C5, Biolegend).

### Quantification

Airway diameters were measured using ImageJ software. The longest and shortest diameter was measured per airway structure and then averaged together. Explant size was measured using a ruler by placing the longest side of the tHLO on the ruler. Ki67+ cells and ECAD+, Ki67+ cells were quantified using a program developed in lab by Kevin Rychel, previously described(Kasputis et al., 2018).

### Experimental replicates and statistics

All experiments were done on at least three (N=3) independent biological samples for each experiment. All error bars represented SEM while the long bar represented the average. Statistical differences were assessed with Prism software using unpaired t-test.

## Supporting information

Table 1 Antibodies

