## Supplementary material for "Human lung organoids develop into adult airway-like structures directed by physico-chemical biomaterial properties": Table 1 Antibodies

**Table 1 Primary and secondary antibody information**

| Primary Antibody | Source | Catalog # | Dilution | Clone |
|---|---|---|---|---|
| Chicken anti-GFP | Aves Lab | GFP-1020 | 1:500 | polyclonal |
| Mouse anti-Acetylated Tubulin (ACTTUB) | Sigma-Aldrich | T7451 | 1:1000 | 6-11B-1 |
| Mouse anti-E-Cadherin (ECAD) | BD Transduction Laboratories | 610181 | 1:500 | 36/E-Cadherin |
| Mouse anti- Human Mitochondria (huMITO) | Millipore | MAB1273 | 1:500 | 113-1 |
| Mouse anti-PLUNC | R&D Systems | MAP1897 | 1:200 | monoclonal |
| Rabbit anti-Cytokeratin5 (CK5) | Abcam | ab53121 | 1:500 | polyclonal |
| Rabbit anti-NKX2.1 | Abcam | ab76013 | 1:200 | EP1584Y |
| Rabbit anti-P63 | Santa Cruz Biotechnology | sc-8344 | 1:200 | H-129 |
| **Secondary Antibody** | **Source** | **Catalog #** | **Dilution** | |
| Donkey anti-chicken 488 | Jackson Immuno | 703-545-155 | 1:500 | |
| Donkey anti-mouse 488 | Jackson Immuno | 715-545-150 | 1:500 | |
| Donkey anti-mouse 647 | Jackson Immuno | 415-605-350 | 1:500 | |
| Donkey anti-mouse Cy3 | Jackson Immuno | 715-165-150 | 1:500 | |
| Donkey anti-rabbit 488 | Jackson Immuno | 711-545-152 | 1:500 | |
| Donkey anti-rabbit Cy3 | Jackson Immuno | 711-165-102 | 1:500 | |
